# Multi-omics data integration approach identifies potential biomarkers for Prostate cancer

**DOI:** 10.1101/2023.01.26.522643

**Authors:** Zedias Chikwambi, Marie Hidjo, Pageneck Chikondowa, Glory Jayeoba, Vincent Aketch, Lawrence Afolabi, Olaitan I. Awe, David Enoma

## Abstract

Prostate cancer (PCa) is one of the most common malignancies, and many studies have shown that PCa has a poor prognosis, which varies across different ethnicities. This variability is caused by genetic diversity. High-throughput omics technologies have identified and shed some light on the mechanisms of its progression and finding new biomarkers. Still, a systems biology approach is needed for a holistic molecular perspective. In this study, we applied a multi-omics approach to data analysis using different publicly available omics data sets from diverse populations to better understand the PCa disease etiology. Our study used multiple omic datasets, which included genomic, transcriptomic and metabolomic datasets, to identify drivers for PCa better. Individual omics datasets were analysed separately based on the standard pipeline for each dataset. Furthermore, we applied a novel multi-omics pathways algorithm to integrate all the individual omics datasets. This algorithm applies the p-values of enriched pathways from unique omics data types, which are then combined using the MiniMax statistic of the PathwayMultiomics tool to prioritise pathways dysregulated in the omics datasets. The single omics result indicated an association between up-regulated genes in RNA-Seq data and the metabolomics data. Glucose and pyruvate are the primary metabolites, and the associated pathways are glycolysis, gluconeogenesis, pyruvate kinase deficiency, and the Warburg effect pathway. From the interim result, the identified genes in RNA-Seq single omics analysis are linked with the significant pathways from the metabolomics analysis. The multi-omics pathway analysis will eventually enable the identification of biomarkers shared amongst these different omics datasets to ease prostate cancer prognosis.

## Introduction

Prostate cancer (PCa) is the most prevalent hormone-dependent oncological disease and the second leading cause of tumour-related deaths among males worldwide (1); (2). Clinically, PCa is characterized by indolent phenotypes, rapid progression and aggressive metastatic disease. Compared to other tumours, PCa progression is slow yet still harms the patient’s long-term health (3). The multifocal and heterogeneity nature of primary PCa is associated with poor prognosis (4), and current clinicopathological indicators do not distinguish well between patients based on their outcomes at the initial stage of the disease (5), as such intervention for patients with metastasis is still urgently needed. Therefore, the efficient identification of the risk level of prostate cancer patients and precise therapeutic targets has always been a critical equation to solve.

Despite its highest genetic diversity, the African population is reported to have a higher prevalence and associated mortality rate of PCa (6) globally._Comparative studies conducted mainly in Africa are crucial to investigate the genetic basis of prostate cancer and its phenotypic adaptation in Africa. Some studies have successfully fine-mapped significant PCa-associated genetic variants using African ancestry datasets. However, the findings could only explain 30– 33% of prostate cancer variability among African individuals (7). A distinct genomic landscape of PCa and genetic polymorphisms associated with specific ethnic groups may lead to different metabolic adaptations that permit tumour cells to increase (6). The lack of adequate genetic reference information from the African genome is one of the significant obstacles in exploring the benefits of precision oncology in the African context. The high ethnic diversity and prevalence of aggressive PCa in the African population represent biomedical research opportunities. The poor understanding of PC pathogenesis has limited the effective clinical management of patients with the PC condition (8). High-throughput omics technologies have revolutionized biomedical research where simultaneous multi-omics data integration helps to unlock hidden biological relationships useful for early diagnosis, prognosis and expedited treatments (9). The recent past has seen rapid development and use of high-throughput technologies, such as genomics, transcriptomics, epigenomics, proteomics and metabolomics, in an attempt to understand PCa (10). In as much as the single omics technologies have identified and shed light on the mechanisms of the tumour progression, subtype and finding new treatment targets, the multisystem and multilevel pathological nature of the disease calls for a holistic molecular perspective of the mechanisms, prospective biomarkers and drug targets and responses. This multi-layered analysis can only be uncovered when a systems biology approach is adopted. Compared to single omics interrogations, multi-omics data integration provides researchers with a greater understanding of the flow of information, from the original cause of the disease (genetic, environmental, or developmental) to the functional consequences or its relevant interactions (11). A practical consideration in multi-omic studies is the correlation of identities of the same objects across omics layers. However, it is possible to infer genetic signatures or phenotypes based on genotypes from different objects (12). The chances of finding a correlation or association between two omics elements that share a common driver or when one factor perturbs the other is very high. Mediation analysis helps to integrate such omics data, treating one as the mediator in the causal mechanism from genotype (SNPs) to phenotype (disease) (13). A study done by Sinha et al. using multi-omics in PCa revealed that proteomics features were significantly more informative for biochemical recurrence prediction than genomic, epigenomics or transcriptomics (14). Integration of transcriptomics and metabolomics reveals significant alterations of several metabolic pathways in PCa (15). Hence, in the current study, we used good quality omics data sets (genomics, transcriptomics and metabolomics) from public databases to better understand tumour progression, subtyping and finding novel biomarkers that potentially address individual variations in drug responses among prostate cancer patients. In addition, we shall also provide a simplified protocol for routine integration of multi-omics data sets to answer biological questions.

## Methods

This study used secondary data on genomics, transcriptomics, and metabolomics from publicly available databases, using a top-down/bottom-up data reduction approach. Quality data was selected with clear quality control standards and accompanying detailed meta-data. Data from each platform were analyzed separately and as an integrated data set to determine the power of the systems biology approach. Because of the multitude of software used for multi-omics studies, the selection of the databases and software/tools was achieved through a screening process followed by performance ranking.

### 1. Genomics (whole genome sequencing) analysis

#### a. Data retrieval and QC

Multiple sequence data were retrieved from SRA using a script-based sra-toolkit to download whole genomic sequence data (PRJNA412953) and converted to fastq format with read 1 and 2 outputted as separate files. Pre-trimming data quality control was performed on the raw reads with MultiQC (43). Afterwards, Trimmomatic-Qc v0.39 (16) was used to trim raw reads with quality score (Q-score) < 10 and < 40 residues in length and remove sequencing adaptors. Confirmation of read quality improvement was then done by post-trimming data quality check with FastQC.

#### b. Read mapping and variant calling

Quality reads were aligned to the reference genome (17) using BWA-MEM (18). The SAM output files from BWA-MEM were converted to BAM format using Samtools (19). Read coverage per position was then calculated using bcftools tools and call variants. Filtering was performed on the SNVs and indels using vcftools, and a variant call file format was generated as the output of the run. snpEff was used for the variant annotation to identify the effects of INDELs and SNVs and were classified based on their functional impact. Finally, the output file was used for SNP Set Enrichment Analysis (SSEA) (20) to provide pathway name, size, enrichment score and nominal p-value associated with prostate samples compared to the control.

### 2. Transcriptomics (RNA-SEQ) analysis

#### a. Data retrieval and quality control

Multiple sequence data were retrieved from SRA using a script-based sra-toolkit to download RNASeq data (PRJNA531736) and converted to fastq format. One hundred eighty-one prostate tissue biopsy cores from Black South African men, 94 with and 87 without pathological evidence for prostate cancer.Pre-trimming data quality control was performed on the raw reads with MultiQC (43). Afterwards, Trimmomatic-Qc v0.39 (16) was used to trim raw reads with Q-score < 10 and < 40 residues in length. Confirmation of read quality improvement was then done by post-trimming data quality check with FastQC.

#### b. Read sequence mapping

The RNA-Seq quality reads were mapped against the Human reference genome (17) using HISAT2 (21). The data was visualised using an integrative genomics viewer (22).

#### c. Differential gene expression analysis

Differential expression analysis was carried out by looking at gene expression values for comparison among samples. The transcript expression level was quantified by aggregating the raw counts of the mapped reads using featureCounts (23). Next, we performed the DGE analysis using DESeq2 (24) between control and prostate cancer samples. Fast Gene Set Enrichment Analysis (fgsea) R package (42) was used to determine the set of genes which were truly enriched in differentially expressed genes. The p-values were then filtered out, which was used as an input file in the multi-omics pathway.

### 3. Metabolomics analysis

Metabolomics data (project ID PR000570) was downloaded from the metabolomics workbench (www.metabolomicsworkbench.org). The metadata and the raw data were inspected to ensure the phenotype metabolites were in rows and columns, respectively. The metabolite profile data was uploaded to the Lilikoi algorithm (25) to standardize the metabolite names from the imputed data to various IDs in databases, including the Kyoto Encyclopedia of Genes and Genomes (KEGG) and Human Metabolome Database (HMDB). In addition, a comprehensive pathway deregulation score (PDS) matrix was generated from the metabolite profiles using the *Pathifier* algorithm (26), and the p-value was subsequently calculated via Chi-square.

### 4. Integrated multi-omics

The goal of our multi-omics integrative analysis was to cluster samples from different omics data sets to identify overlapping biomarkers. The multi-omics data analysis comprised mainly three broad categories, Regression/Association-based Methods, Clustering-based Methods, and Network-based Methods. The p-values of up-regulated genes from the transcriptomics pathway, p-values from the lilikoi pathway for metabolomics and p-values of the enrichment score from whole genome sequence analysis were uploaded into the Pathway multi-omics tool’s MiniMax matrix (27), which sort each omics p-value and output the minimal p-value that links with pathways in at least two of the omics.

## Results

### 1. RNA-Seq Analysis Results

The differentially expressed genes from the RNA-seq data are shown in figure 5A. The genes in normal prostate tissue have a uniform expression. However, the genes in prostate tumours have differentially expressed genes between the two phenotypes and within the prostate tumour samples themselves. Numerous studies have also shown that some genes are highly expressed in prostate cancer tumours (28–30). The volcano plot in figure 5B shows the list of genes highly expressed in the prostate tumour, and GLYATL1, RCC1, AMACR, UBE2E3-DT and RAB3B are the up-regulated genes and EFNB1 CSTA, SPON1, and GLIST are down-regulated. The cell cycle pathway from the Kyoto Encyclopedia of Genes and Genome in figure 5C shows some of the genes from the cell cycle that are highly expressed, where some genes that are involved in DNA damage checkpoint (ATMATR) are lowly expressed, and some genes involved in DNA biosynthesis are highly expressed (ORC, MCM).

**Figure 1:**
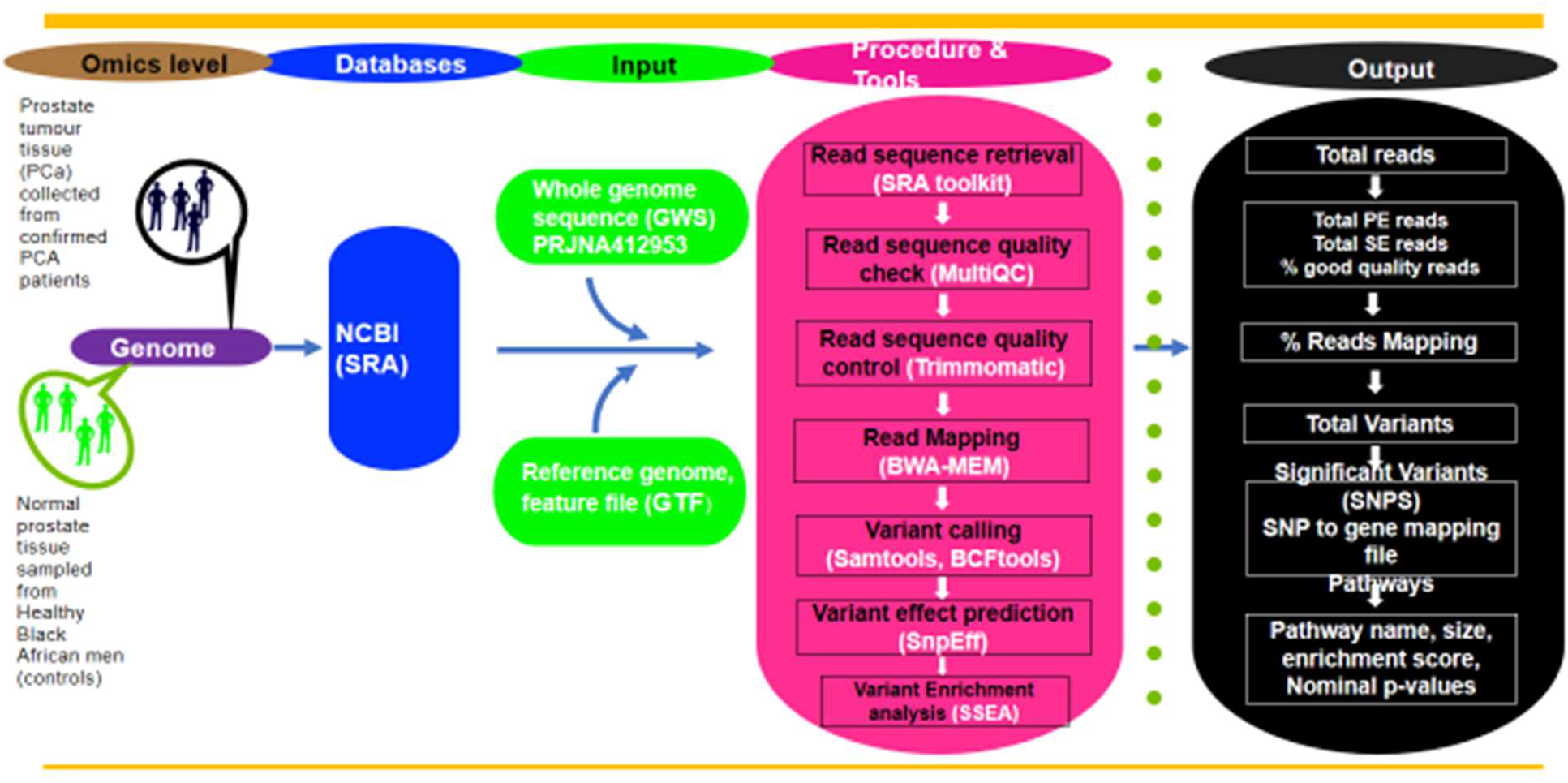
The workflow of genomics analysis. It uses WGS data as an input file consisting of 7 procedures: Sequence retrieval, Quality check, quality control, Mapping, Variant Calling, Variant Prediction and Variant Enrichment Analysis.

**Figure 2:**
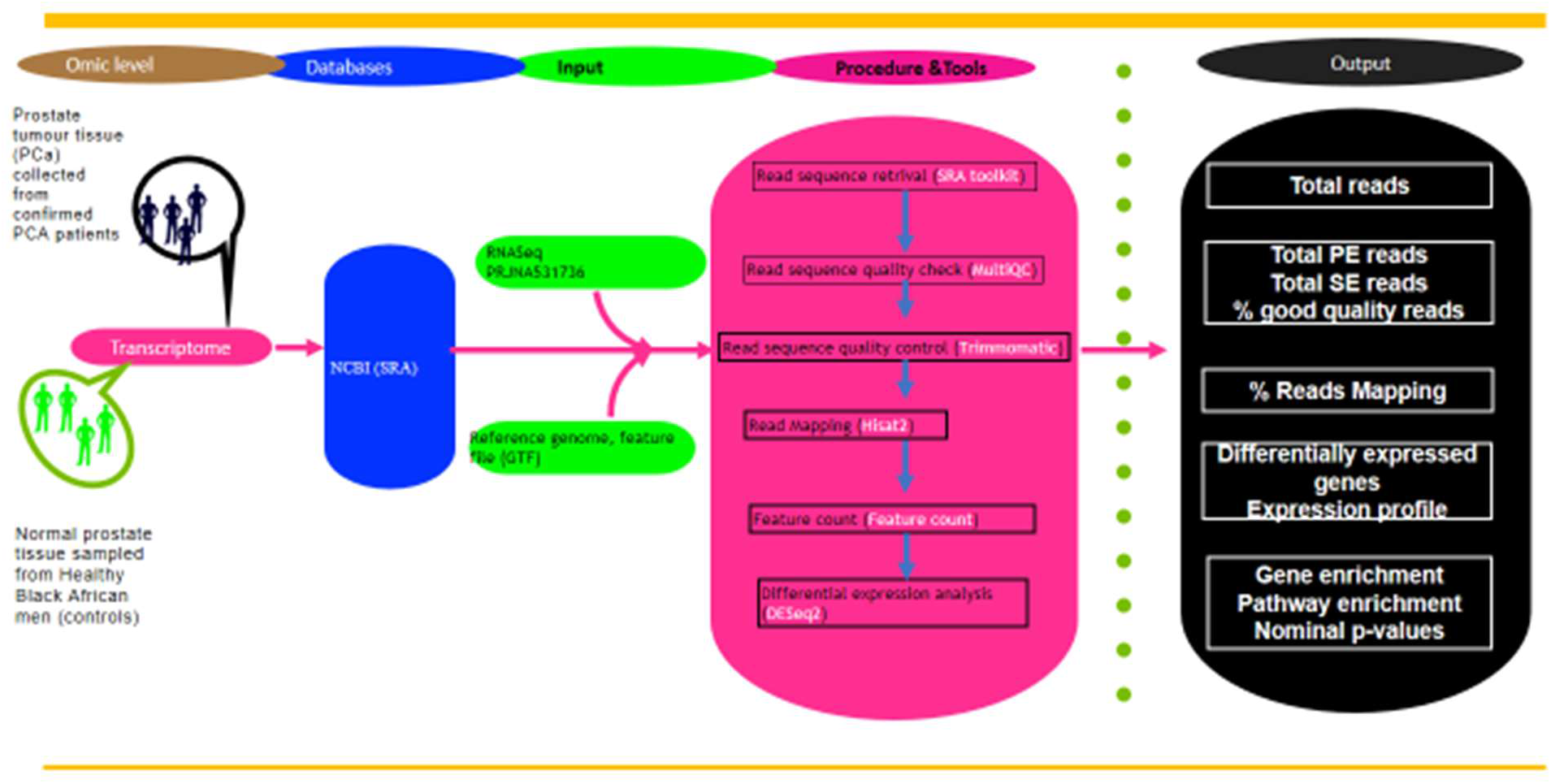
The workflow of transcriptomics analysis. It uses RNA-Seq data as an input file consisting of 6 procedures: Sequence retrieval, Quality check, Quality control, Mapping, Feature Counts and Differential Gene expression analysis.

**Figure 3:**
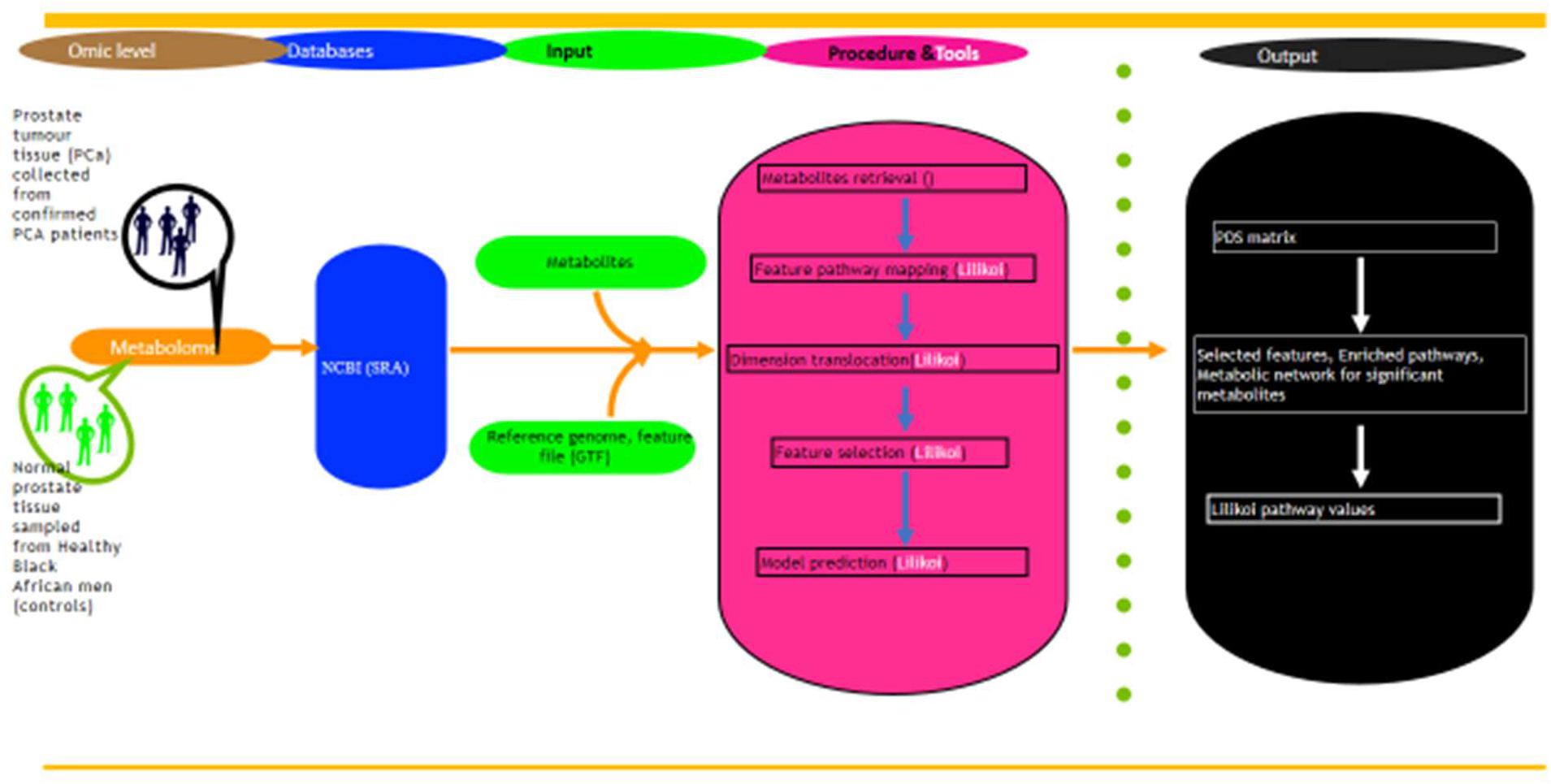
The workflow of metabolomics analysis. It uses Lilikoi v2.0 R package composed of a feature mapper, preprocessing, dimension transformer, exploratory analysis, and pathway analysis. Input data require a metabolomics data matrix and 1 column of a categorical variable to specify the case/control status for each subject. The feature mapping converts metabolite names to standardized metabolic IDs and transforms them into pathway names.

**Figure 4:**
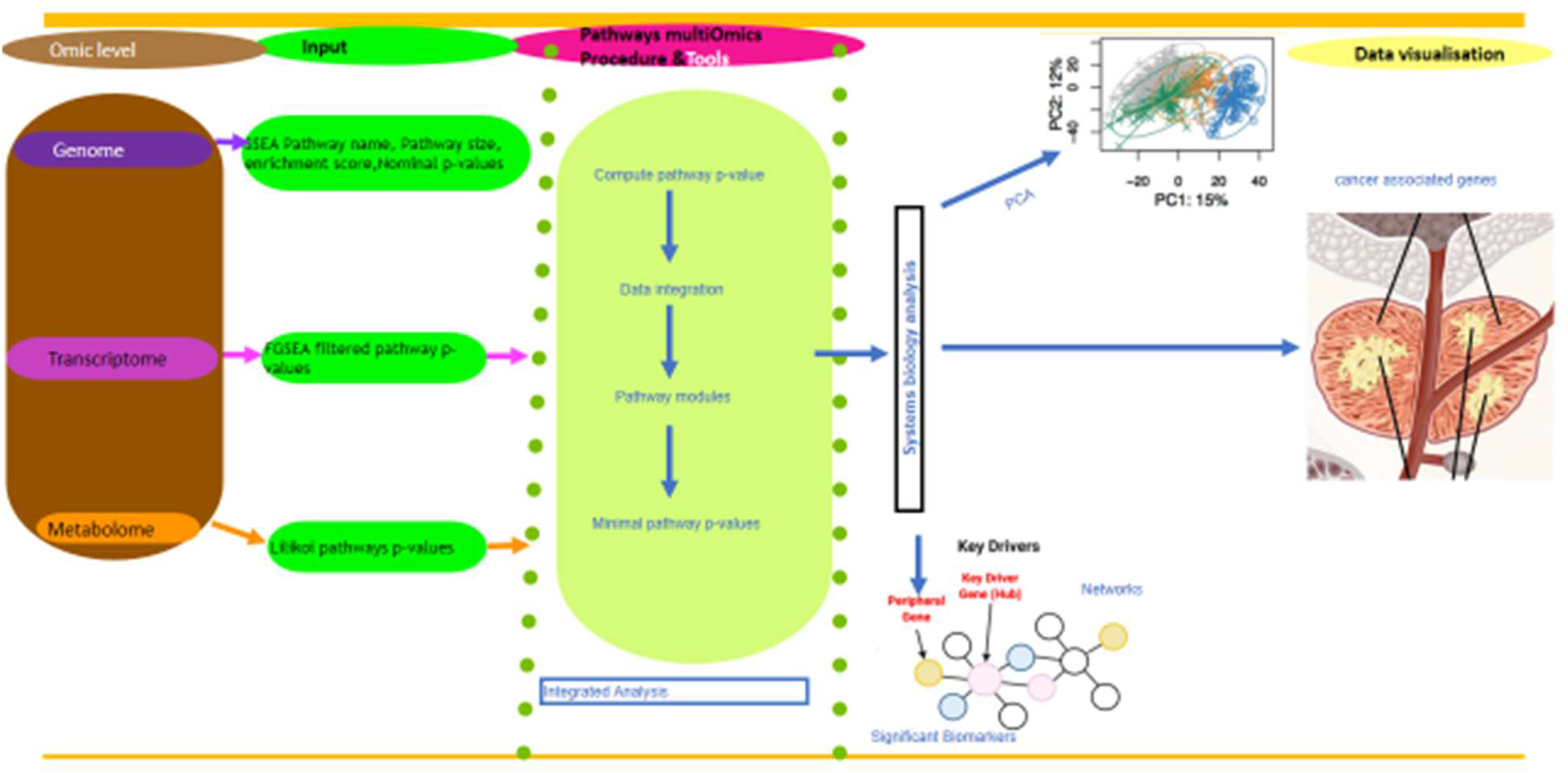
The workflow of multi-omics analysis. It comprises mainly of three broad categories, the p-values of the enrichment score analysis from the whole genome sequence pathway, p-values of up-regulated genes from transcriptomics and p-values from the lilikoi pathway for metabolomics is used as an input into the MiniMax matrix, and this produces common pathways from at least two of the omics.

**Figure 5:**
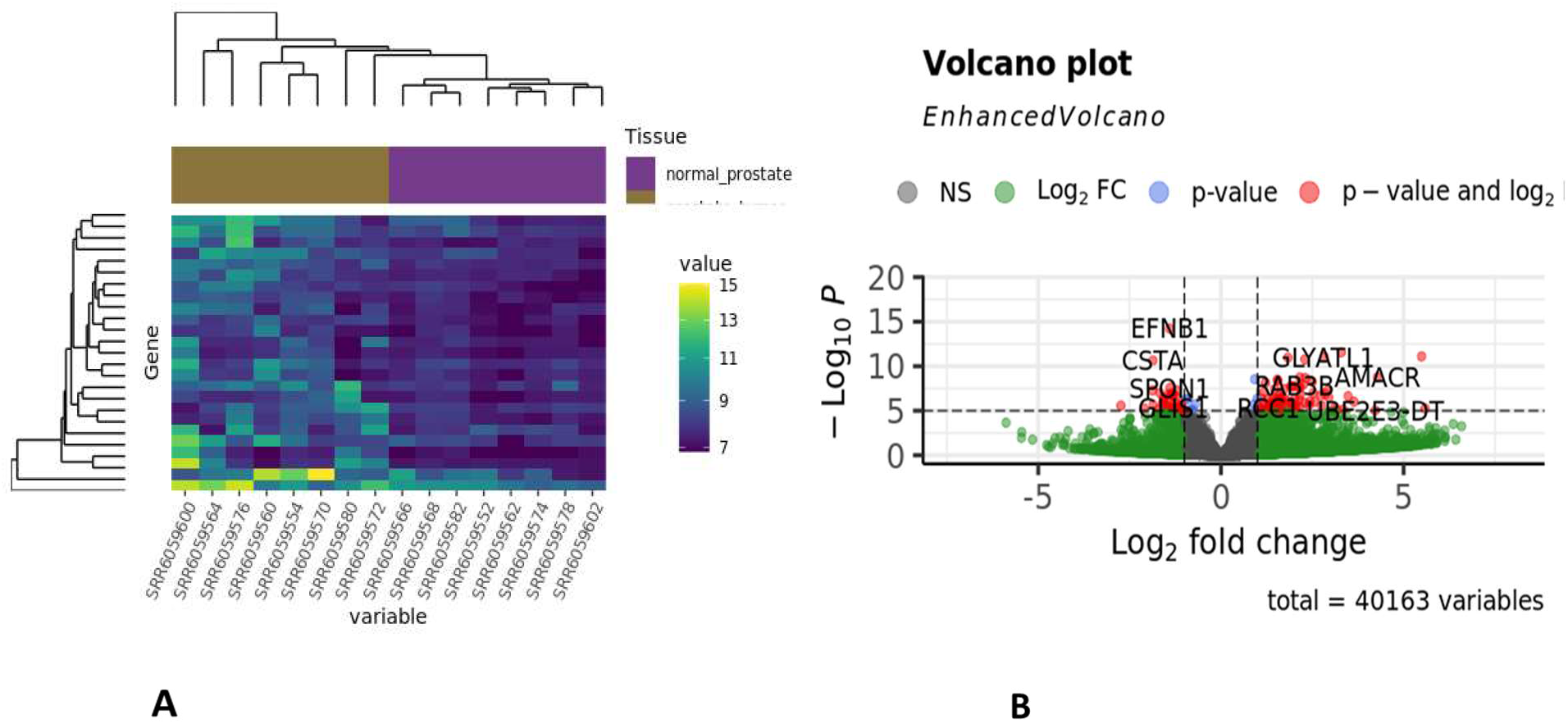
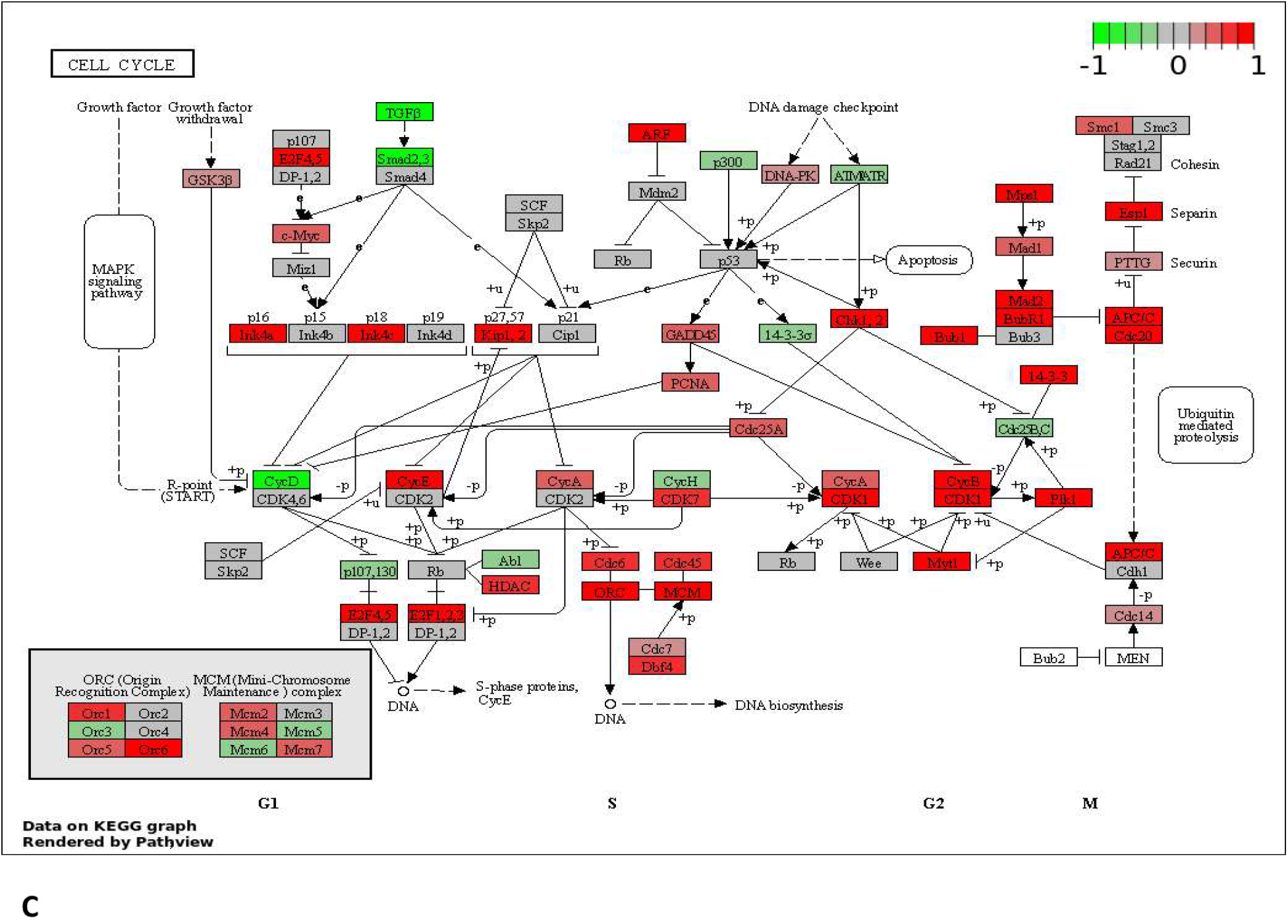
RNA-Seq single omics results. **(A) Clustering and genes distance.** Purple indicates normal tissue with the genes having similar expression, and Tortilla shows the prostate tumour samples with the genes being expressed differently than the normal. **(B) Differentially Expressed Genes**. The red dots at the right represent genes that are highly expressed, and those at the left are genes that are downregulated, while the grey dots in the middle have normal expression. **(C) Cell cycle pathway associated with prostate cancer**. The overlay of RNA-Seq data in the Kyoto Encyclopedia of Genes and Genome showed different pathways involved in the prostate transcript. The genes in red are up-regulated, genes in green are downregulated, and genes in grey have normal regulation

### 2. Metabolomics Analysis Results

From the metabolomics dataset, eleven pathways were identified as feature pathways in the prostate tissue (Figure 6B). Galactosemia, Galactose Metabolism, Carbon Metabolism, Biosynthesis of Amino Acids, Amino Sugar and Nucleotide Sugar Metabolism, Alanine, Aspartate, and Glutamate Metabolism are the pathways most relevant to prostate tissue with the highest information gain score > 0.010. In figure 6B, the major metabolites associated with these top pathways are Geranyl diphosphate, Fructose, Glucose, Mannose, and Pyruvate. The pathways and their corresponding metabolites were visualised using the partite graph at (*P* < 0.05), where the cyan nodes are metabolites and the yellow nodes represent pathway features. The metabolic pathways are associated with the largest number of metabolites (Figure 6C). Among them, dihydroxyacetone phosphate, pyruvate, and 6-phosphoglucose acid have the most weight on edge. Many pathways related to amino acid synthesis and metabolism were also highlighted.

Metabolomics

**Figure 6:**
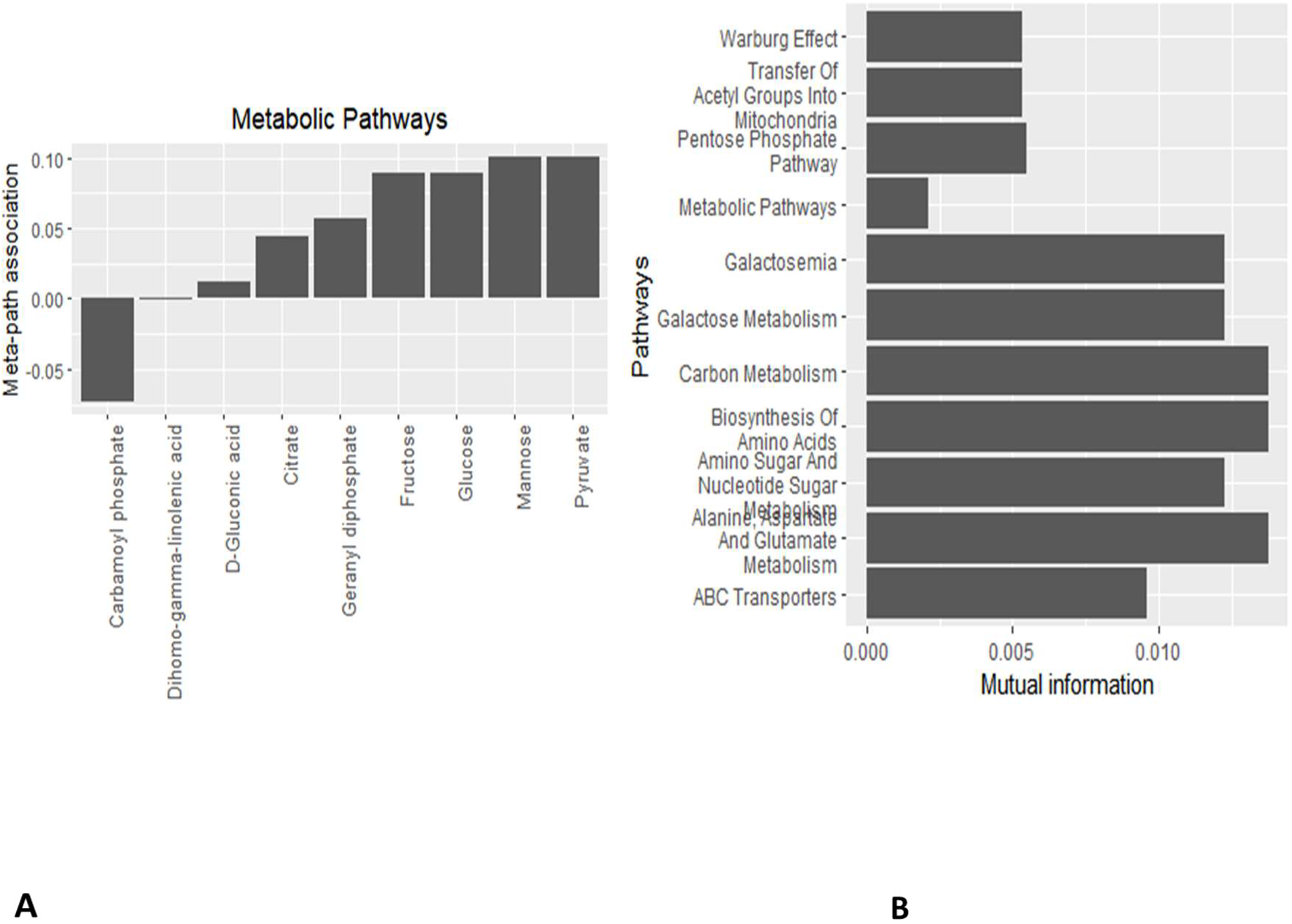
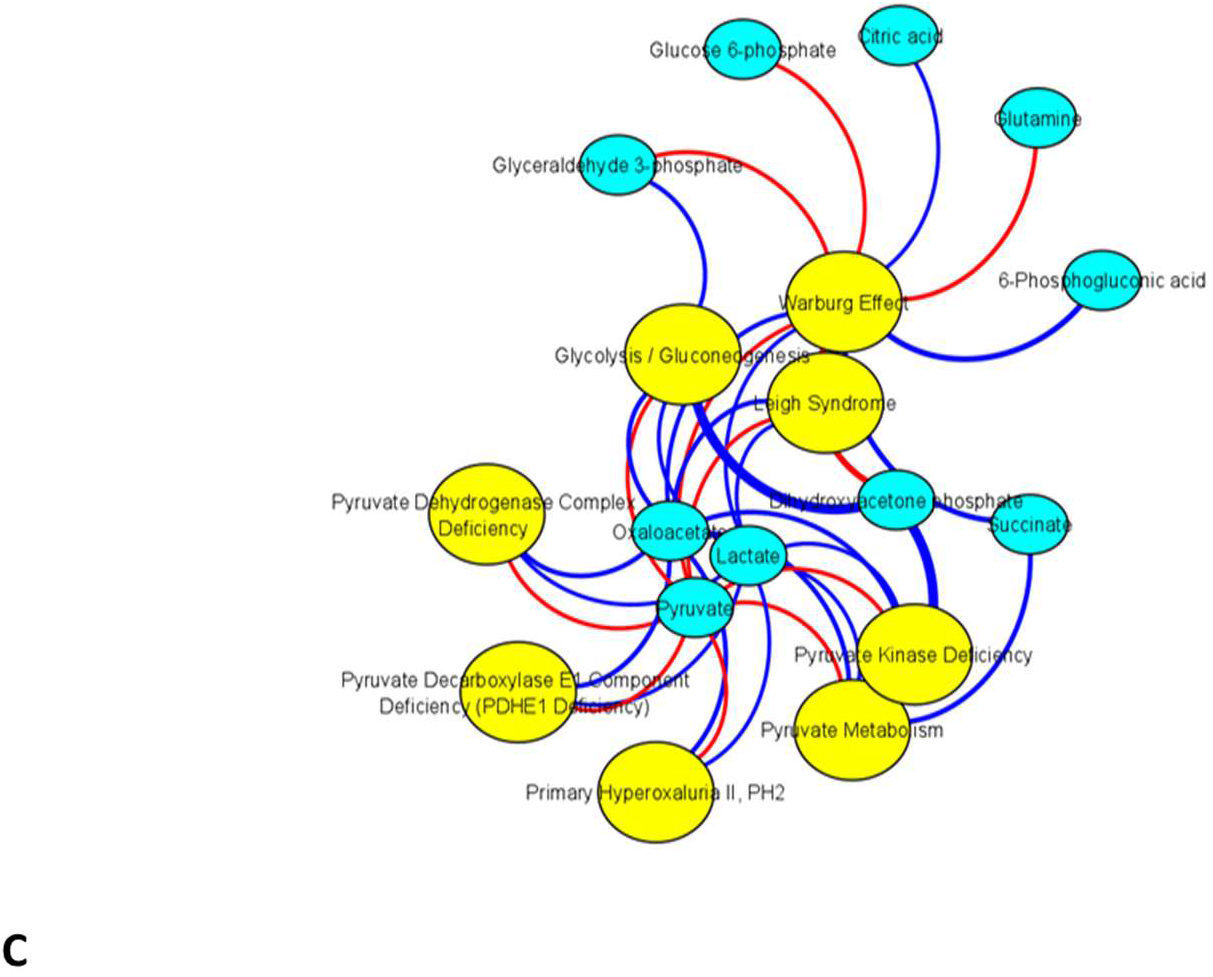
Metabolomics single omics metabolite-pathway relationship results. (A) Metabolites Relationship Bar plots. It shows the relationship between the Citrate, Geranyl diphosphate, Fructose, Glucose, Mannose, and Pyruvate metabolism pathways. (B) Measured Pathway Features. In the measured pathway features, the x-axis represents an information gain score that measures the importance of the pathways, and the y-axis displays the names of pathways. (C) Bipartite plot with eight pathways and their corresponding metabolites. The cyan nodes indicate the metabolites, and the yellow nodes indicate pathways. The red edges are negative associations, the blue edges are positive associations, and the thicker edges indicate higher levels of association.

### 3. Integrative Mult-Omics approach

The Minimax pathway-based approach for integrative analysis of multi-omics data with categorical, continuous, or survival outcome variables. The input of P athwayMultiomics is pathway p -values for individual omics data types, which are then integrated using a novel MiniMax statistic, to prioritize pathways dysregulated in multiple types of omics datasets.

## Discussion

In this multi-omics study, we used publicly available data on genomics, transcriptomics and metabolomics from PCa patients conducted on African ancestry. We observed a similar metabolic pathway in the RNA-Seq and metabolic data sets from African ancestry. Studies have shown that the metabolism patterns of cancer cells are very different from tumour cells, but they both use similar pathways, cell growth differentiation and maintenance (31). There are numerous molecular alterations during prostate cancer progression (32); for example, from our analysis, the glycolytic pathway is highly regulated, which provides the building block for synthesising precursors needed for cell growth. The prostate cancer metabolic analysis in figure 1 shows the up-regulated pathways and their corresponding metabolites from the African American population. Glucose and pyruvate are the major metabolites associated with glycolysis, gluconeogenesis, pyruvate kinase deficiency, and the Warburg effect pathway. One of the major hallmarks of cancer growth is the ability to sustain its proliferation and tumour angiogenesis (33) (31). Pyruvate is an important metabolite from glycolysis, which links many metabolic pathways together (34). The upregulation of glycolytic pathways is most likely associated with the high energy needed for prostate cancer cells to support their fast proliferation.

From the up-regulated cell cycle pathway, E2 factor (E2F), Checkpoint kinase 1 (Chk1), and Minichromosome Maintenance Complex (MCM) are the up-regulated gene and Transforming Growth Factor (TGFR) is down-regulated. The up-regulated genes are majorly used by cancer cells to promote tumour growth, cell proliferation and control of cell cycle DNA replication. The downregulated gene is involved in inhibiting hepatocytes and epithelial cell growth, and its downregulation indicates that prostate cells are growing without control. RAB3B genes have been previously shown to be up-regulated in prostate cancer cells to promote cell proliferation via NKX3-1 and AR regulation pathways and networks (35) (36). The RAB gene is a family of GTPases that regulates cancer progression, enhancing cancer progression, and it has been linked with poor prognosis (37). The mechanism of up-regulated RAB38 gene is most likely due to the increase in mitochondrial respiration. The metabolic pathways associated with pyruvate metabolites are channelled towards oxidative phosphorylation to enhance cell growth. The highly expressed N-acytransferase (GLYATL1) in the present study analysis agrees with other studies showing increased gene expression in prostate cancer cells (38–40). The upregulation of GLYATL1’s gene has been associated with the early-stage progression of prostate cancer. The knockdown of this gene has shown some involvement in the glycolytic pathway, where it inhibits cell proliferation (38). This explains why high metabolites are channelled towards the glycolysis pathway in our metabolic pathway analysis from the metabolomics data. The Alpha-methyl acyl-CoA racemase (AMACR) is an enzyme involved in the biosynthesis of fatty acids, and it has been shown to be highly expressed in some cancer cells (41). The overexpression of the AMACR gene in our RNA data suggests a link with prostate cancer. However, some studies have indicated that the expression of the AMACR gene in a tumour is not enough to be used as a diagnostic tool but can be combined with other biomarkers.

### Conclusions and next steps

It is now known and well-acknowledged that identifying biomarkers that can help in the early detection, prognosis and treatment of prostate cancer will benefit and save lives, especially those of African ancestry. Our holistic approach identified some common biomarkers in at least two omics data sets, as discussed earlier in the multi-omics pipeline. To achieve the current efforts of personalised therapy in treating PCa, a complete holistic approach that will include more epigenetic and proteomic data to identify more biomarkers that might strengthen the present ones is needed. However, a wide range of genetic consortia efforts is still necessary to interrogate the genomes of the African population by pooling efforts and resources specifically to address prostate cancer in Africa.

## Availability and Requirements

Multi-Omic Data Analytics Integration in Prostate Cancer https://github.com/omicscodeathon/cancer_prostate

Programming language: Bash, R 3.5, Python.

Other requirements: Python3.6 or higher, Java JDK, Chrome, Firefox, and Safari web browser.

License: MIT

Any restrictions to use by non-academics: None.

## Availability of data and materials

The dataset analysed during the current study as a case study is publicly available at NCBI Sequence Read Archive (https://www.ncbi.nlm.nih.gov/sra) with Project IDs PRJNA412953 & PRJNA531736. The metabolomics data PR000570 on Metabolomics workbench (www.metabolomicsworkbench.org) The Project repository, which also includes the entire code and other requirements, can be downloaded from https://github.com/omicscodeathon/cancer_prostate .

## Abbreviations

AMACR: Alpha-methylacyl-CoA racemase
CLI: Command-Line Interface
FDR: False Discovery Rate
GUI: Graphical User Interface
E2F: E2 factor
Chk1: Checkpoint kinase 1
MCM: Minichromosome Maintenance Complex
TGFR: Transforming Growth Factor
RAB3B: Ras-related protein Rab-3B

## Acknowledgements

The authors thank the National Institutes of Health (NIH) Office of Data Science Strategy (ODSS) and the National Center for Biotechnology Information (NCBI) for their immense support before and during the April 2022 Omics codeathon organised in collaboration with the African Society for Bioinformatics and Computational Biology (ASBCB). The authors acknowledge the NIH Library Editing Service, for manuscript editing assistance.

